# Forebrain neuronal SMC3 regulates body weight and metabolic health partially through regulation of hypothalamic Melanocortin 4 receptor

**DOI:** 10.1101/2024.04.07.588459

**Authors:** Natalia Saleev, Dmitriy Getselter, Joanna Bartman, Roee Gutman, Asaf Marco, Evan Elliott

## Abstract

SMC3 is a major component of the cohesin complex that regulates higher-order chromatin organization and gene expression. Mutations in SMC3 gene are found in patients with Cornelia de Lange syndrome (CdLs). This syndrome is characterized by intellectual disabilities, behavioral patterns as self-injury, as well as metabolic dysregulation. Nonetheless, little is known about the role of neuronal SMC3 in gene expression and physiology in adulthood. This study determined the role of SMC3 in adulthood brain, by knocking out Smc3 specifically in adulthood forebrain excitatory neurons. Excitatory conditional neuron-specific SMC3 knockout (cKO) mice displayed a very strong metabolic phenotype in both male and female mice, including a robust overweight phenotype, loss of muscle mass, increased food consumption, lower respiratory exchange ratio, lower energy expenditure and hormonal changes. The hypothalamus displayed dysregulated neuronal morphology and associated transcriptional abnormalities in RNA-seq analysis across various cellular pathways, including decrease of Melanocortin 4 receptor (*Mc4r*) expression level, a pivotal regulator of appetite and metabolism. Correspondingly, ChIP-seq analysis revealed genome-wide alterations in the binding dynamics of SMC3 of the cKO animals, including Mc4r associated regions. Notably, a significant correlation emerged between multiple sites exhibiting a marked decrease in binding and downregulated genes. The administration of Setmelanotide, an MC4r agonist, to cKO group resulted in a notable reduction in both weight and food consumption in these mice. Therefore, we have identified specific and reversable metabolic parameters that are regulated by neuronal Smc3 in adulthood.

## Introduction

SMC3 (Structural maintenance of chromosomes protein 3) is a major component of the cohesin protein complex that regulates higher-order chromatin organization and gene expression^1,2^. Although the cohesin complex is best known for its function in cohesion of sister chromatids during the cell cycle, cohesin has been implicated in coordination of transcriptional programs by promoting or preventing the interaction between desired regions, such as promotors, enhancers and inactivator regions^3^. SMC3 is ubiquitously expressed in all cell types and is considered mandatory for proper cellular development.

De novo mutations in multiple members of the cohesin complex, including *SMC3* gene, are associated with the developmental disorder Cornelia de Lange syndrome (CdLs)^4,5^. This syndrome is characterized by intellectual disabilities, behavioral dysregulation, such as self-injury, as well as metabolic dysregulation^6^. Children born with CdLs often have low birthweight, with poor ability to gain weight^7^. However, a clinical report of Italian adults with CdLs reported that 46.5% of screened people were overweight. A separate study from USA found that 18% of their cohort of adults with CdLs were overweight^8^. Therefore, while CdLs is often associated with undergrowth in development, there is often an opposite phenotype by adulthood.

Homozygous knockout of *Smc3* leads to embryonic lethality, thus at least one allele is needed to enable embryonic development^9^. Heterozygous loss of *Smc3* in developing mice brain and tau-expressing neuronal cells increased dendritic complexity in cortical layers, together with a behavioral phenotype including anxiety^10,11^. An in-vitro study found that cohesin complex is necessary for proper gene transcription in post-mitotic neurons^12^. However, the role of SMC3 in post developmental maintenance of neuronal function in an animal has not been determined.

To understand the function of SMC3 in the adulthood brain, we established a neuron-specific knockout of *Smc3* specifically in adulthood in mice. Adulthood neuron-specific knockout of *Smc3* induced a robust metabolic phenotype, including increased food consumption, weight gain, and dysregulated hormonal balance, in association with morphological and transcriptional changes in hypothalamus. These changes were attenuated with Setmelanotide, an agonist of melanocortin 4 receptors. The results indicate that neuronal adulthood SMC3 has a particularly important role in regulating proper metabolism during adulthood.

## Results

### Knockout of *Smc3* gene in excitatory neurons

To understand the role of neuronal SMC3 protein in adult brain, we developed tamoxifen-inducible knockout mice for *Smc3* gene in excitatory neurons, by using Camk2a-cre/ER^T^2 mice that were crossed with floxed *Smc3* (on exon 4) mice^13^. Camk2a promoter promotes expression specifically in forebrain excitatory neurons, and Camk2a-cre/ER^T^2 mice have been extensively used to study the function of specific genes in these cell types^14–16^. Tamoxifen was given by gavage at eight weeks of age which is associated with adulthood in mice. Immunostaining of hippocampus confirmed the knockout of *Smc3* 4 weeks after tamoxifen treatment (Fig. 1A, B). Furthermore, we see that all mice survive the lack of *Smc3* for at least 15 weeks after tamoxifen treatment (Fig. 1C, D). In addition, TUNEL staining determined that there is no difference in apoptotic cells in the hippocampus between the two experimental groups (Fig. 1E, F). Therefore, SMC3 is not necessary for neuronal survival during adulthood.

**Figure 1.**
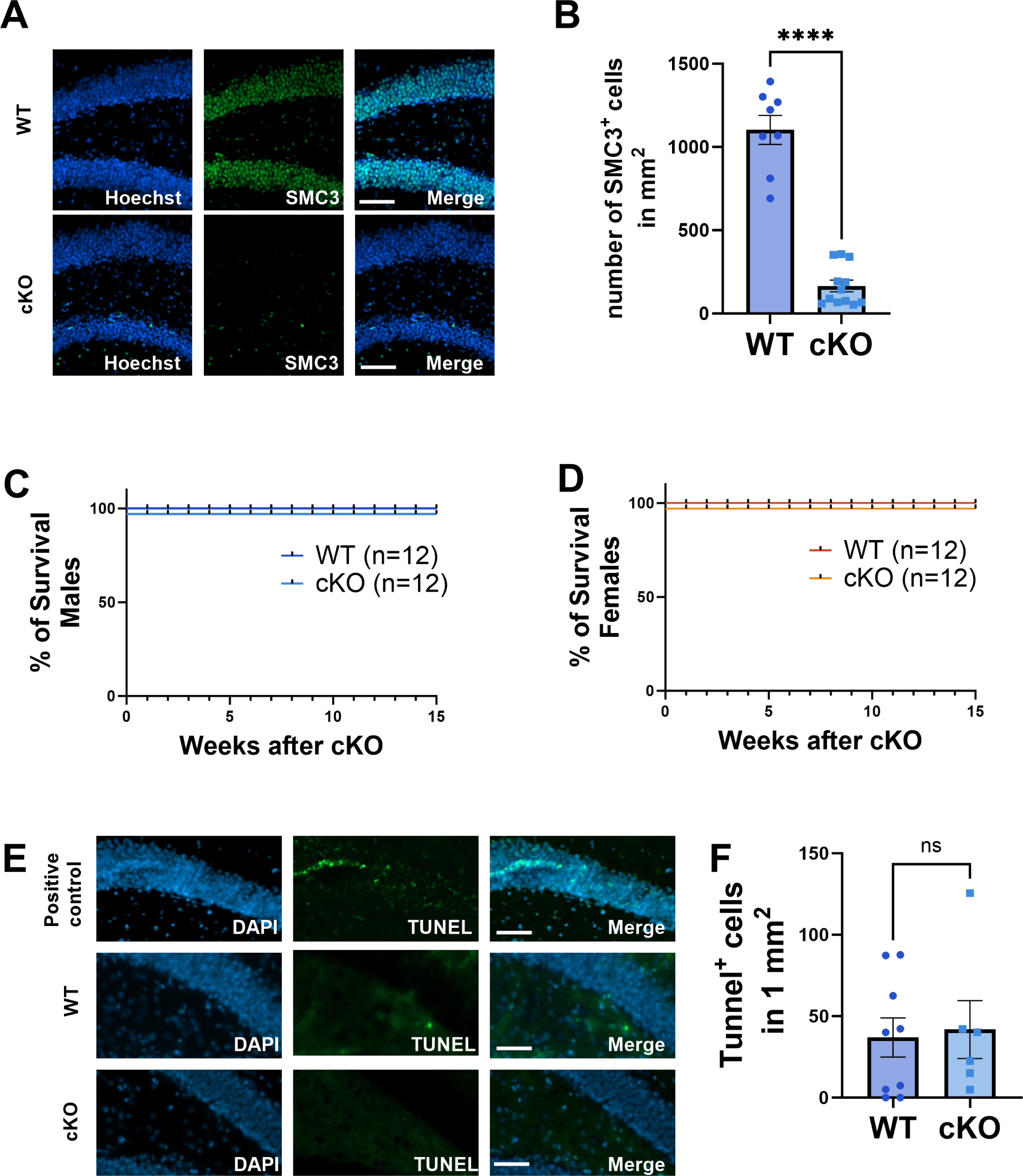
Knockout of *Smc3* gene in excitatory neurons during adulthood. (**A)** Representative pictures of hippocampus in wild type and *Smc3* knockout mice4 weeks after tamoxifen treatment.; Scale bars present 100*μm*. **(B)** quantification of SMC3 positive cells in DG of both groups (two-sided *t*-test WT, n=8; cKO, n=12; ****P<0.0001). **(C-D)** Kaplan-Meier male and female survival curves. The survival of mice from tamoxifen treatment for 15 weeks shows that lack of SMC3 does not cues lethality. **(E)** TUNEL staining for cell apoptosis at 4 weeks after tamoxifen treatment. scale bars represent 200*μm*. **(F)** Quantification of apoptosis cells shows that there is no difference between *Smc3* cKO and WT mice (two-sided *t*-test WT, n=9; cKO, n= 6).

### Metabolic effect as result of *Smc3* gene cKO in adult brain

Mice were observed for any noticeable physical abnormalities. Surprisingly, *Smc3* cKO mice were visibly larger than wild type mice by week 5 after tamoxifen treatment (Figure 2A). Mice were measured as several time points, and significant weight different between genotypes developed by day 34 after tamoxifen treatment in both male and female mice (Figure 2B, C). In parallel, *Smc3* cKO mice displayed increased food consumption (Figure 2D, E). Nuclear Magnetic Resonance NMR spectroscopy, 5 weeks after tamoxifen treatment, revealed that fat tissue was increased, and lean body tissue was decreased in *Smc3* cKO mice (Figure 2F—I). Interestingly, we further found that the same pattern of changing in body composition was already present at day 28 after tamoxifen treatment (Supplementary Figure 1), before the weight of wild type and *Smc3* cKO mice were significantly different.

**Figure 2.**
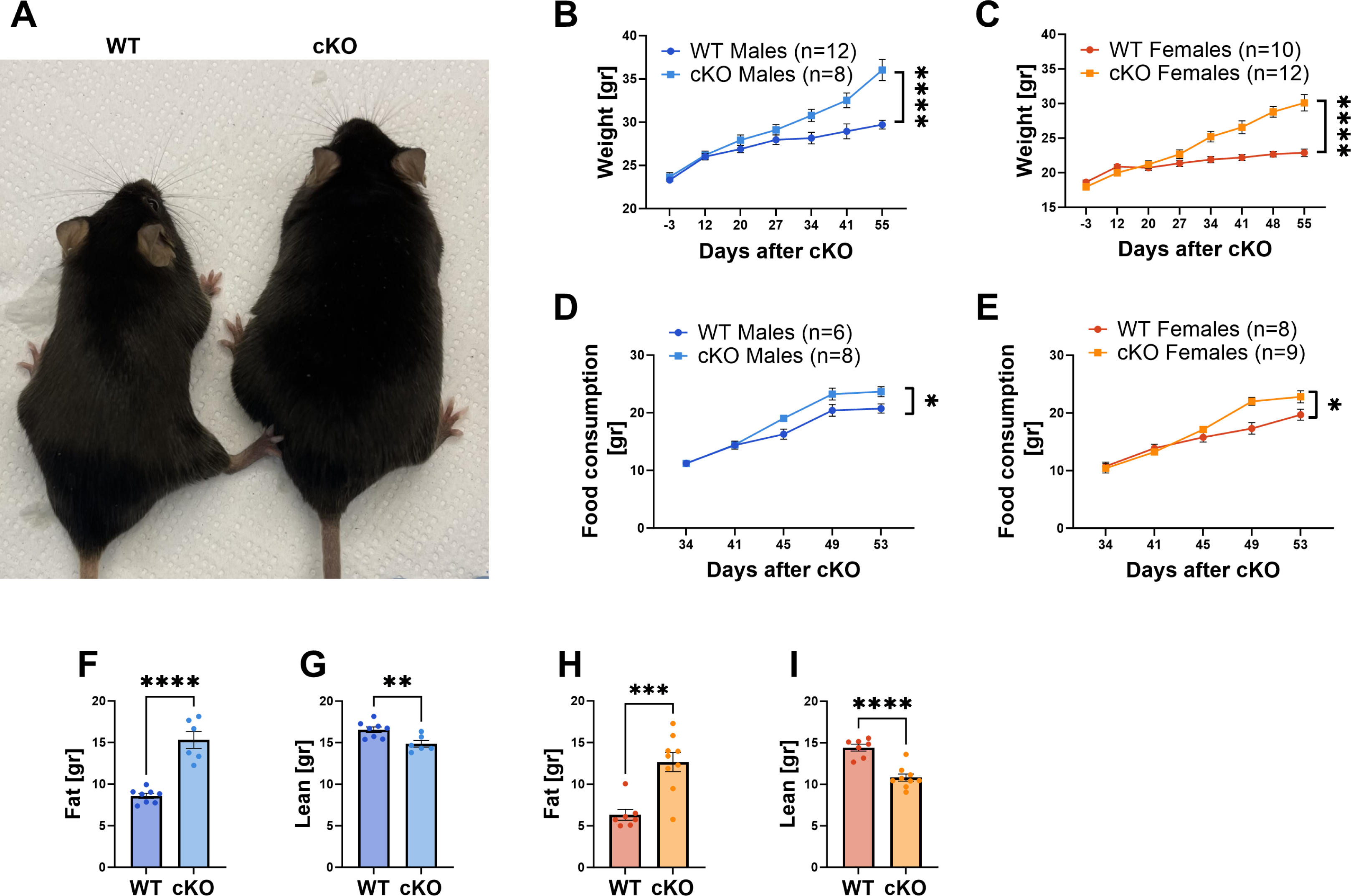
Metabolic effect as result of *Smc3* gene cKO in adult brain. **(A)** Representative picture of male mice at week 12 after tamoxifen treatment. **(B)** Male body weight measured show significantly increase from 34^th^ day onwards (WT, n=12; cKO, n=8; F_6,128_=8.994; ****P<0.0001, Two-Way ANOVA interaction test). **(C)** Female body weight measured shows significant increase from 34^th^ day onwards (WT, n=10; cKO, n=12; F_7,145_=28.95; ****P<0.0001, Two-Way ANOVA interaction test). **(D)** Food consumption by male mice measured between cKO and WT group over time (WT, n=6; cKO, n=8; F_4,70_=3.557; *P=0.0106, Two-Way ANOVA interaction test). **(E)** Food consumption by female mice measured between cKO and WT group over time (WT, n=8; cKO, n=9; F_4,77_=2.719; *P=0.0356, Two-Way ANOVA interaction test). **(F-G)** Body composition in male mice of Fat and Lean tissue measured using Mini-spec system at day 40 after tamoxifen treatment (two-sided *t*-test WT, n=8; cKO, n=6; ****P<0.0001; **P=0.0069). **(H-I)** Body composition in female mice of Fat and Lean tissue measured using Mini-spec system at day 40 after tamoxifen treatment (two-sided *t*-test WT, n=7; cKO, n=9; ****P<0.0001; ***P=0.0006).

To gain more insight into metabolic differences between phenotypes, mice were observed in metabolic chambers. *Smc3* cKO mice displayed a lower respiratory exchange ratio (Fig. 3A) and heat production (Fig. 3B). In addition, *Smc3* cKO mice did not display the typical circadian rhythms that are present in the wild type mice. However, there were no differences between phenotypes in total distance moved inside chambers (Fig. 3C). Therefore, the metabolic changes are not due to changes in movement.

**Figure 3.**
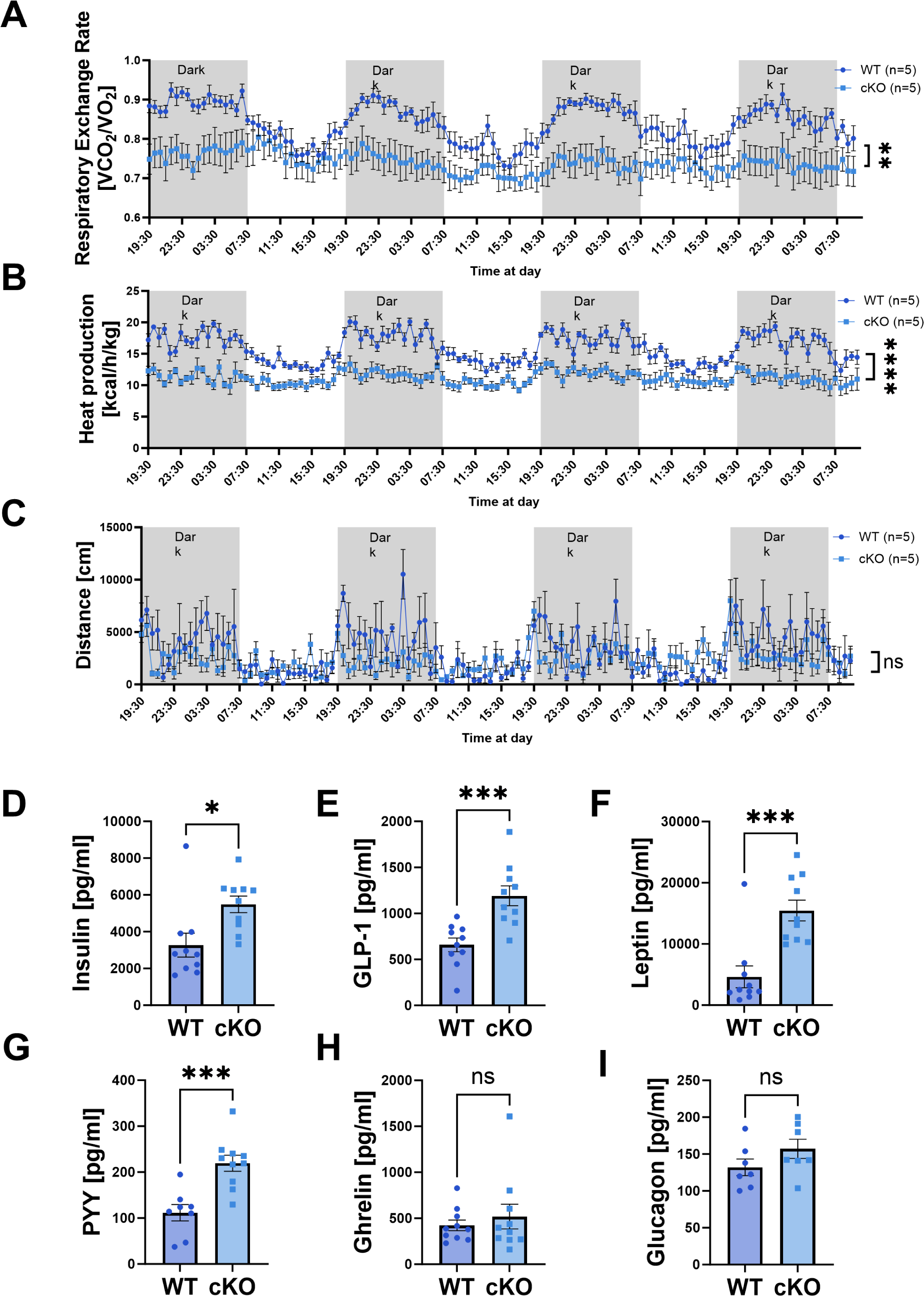
Metabolic differences in *Smc3* cKO mice. **(A-C)** Metabolic parameters as recorded in Metabolic chambers. Respiratory exchange rate [RER] (A), heat production (B) and distance moved at the chamber (C). Significant reduce between WT and cKO at main group-effect were indicated at RER and heat production, but not at moved distance (WT, n=5; cKO, n=5; ****P<0.0001; **P=0.0024; not significant (ns) P>0.05, Two-Way ANOVA main group effect). **(D-I)** Hormone analysis of male mice serum six weeks 6 after tamoxifen treatment. (H-I) when cKO compared to WT; Insulin, GLP-1 (glucagon like peptide 1), Leptin, and PYY (Peptide YY or Peptide tyrosine tyrosine) (D-G) were indicated as significantly different when compared cKO to WT mice (two-sided *t*-test n=10; ***P<0.001; *P<0.05; not significant (ns) P>0.05).

In order to understand hormonal differences that may explain changes in food consumption and weight gain, relevant hormonal levels were determined. At 6 weeks after tamoxifen treatment, PYY, GLP-1, leptin, and insulin, but not ghrelin and glucogen, were significantly increased in cKO group (Fig 3D-G) (Fig. H, I). By 9 weeks after tamoxifen treatment, glucagon was also significantly increased in the *Smc3* cKO animals (Supplementary Figure 2). These findings are consistent with a physiological attempt to decrease food consumption and attenuate the increase of body weight. Therefore, increased weight is not due to a dysregulation of peripheral hormones.

### Changes in hypothalamic dendritic branching and gene transcriptions as result of *Smc3* knock out

Considering the metabolic phenotype, and the fact that the hormonal response is consistent with a peripheral attempt to attenuate food consumption and weight gain, we hypothesized that the hypothalamus may be dysregulated in the *Smc3* cKO mice. Using immunohistochemistry, we verified the decrease of Smc3 in the paraventricular nucelus of the hypothalamus in *Smc3* cKO mice (Supplementary Figure 3). Initial characterization of the hypothalamus was performed by Golgi staining to determine neuronal morphology. Dendritic branching was significantly decreased in the hypothalamus of *Smc3* cKO mice in both the male and female cohorts (Fig. 4).

**Figure 4.**
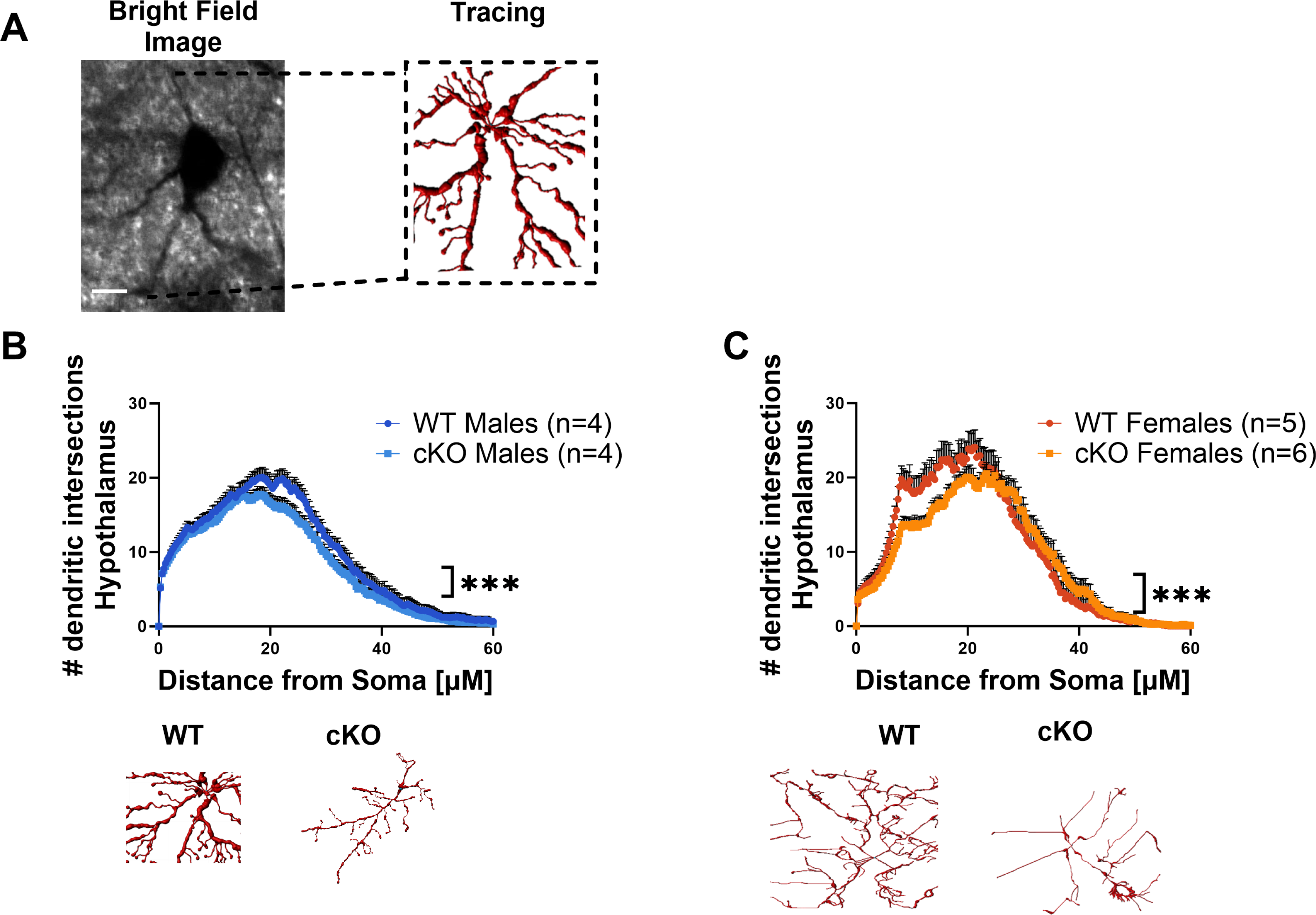
Changes in dendritic branching as result of *Smc3* knock out. Tracing hypothalamic neurons in both sexes, made by Golgi staining that served as template for 3D reconstruction. **(A)** Representative reconstructed hypothalamic neuron. Scale bar present 10 *μm*. **(B)** Males cKO group show decreased dendritic intersections in hypothalamus 11 week after cKO induced (n=4; WT R^2^=0.559; cKO R^2^= 0.5809; CI=99%; ***P<0.001, Non-linear regression). **(C)** Females cKO group show decreased dendritic intersections in hypothalamus 11 week after cKO induced (WT, n=5; R^2^=0.5525; cKO, n=6; R^2^= 0.5449; CI=99%; ***P<0.001, Non-linear regression).

To understand if Smc3 is regulating the hypothalamic transcriptome, whole transcriptomic analysis was performed on RNA extracted from the hypothalamus of wild type and Smc3 cKO knockout mice. Analysis of differentially expressed genes (DEGs) identified 1072 genes downregulated and 532 upregulated genes (Fig. 5A, B, Supplementary Table 1) in the *Smc3* cKO mice. Gene ontology analysis found that upregulated genes were enriched in pathways involved in mitochondrial function. Interestingly, downregulated DEGs exhibited enrichment for genes associated with functions in dendrites, synapses, and the extracellular matrix (Fig. 5C).

**Figure 5.**
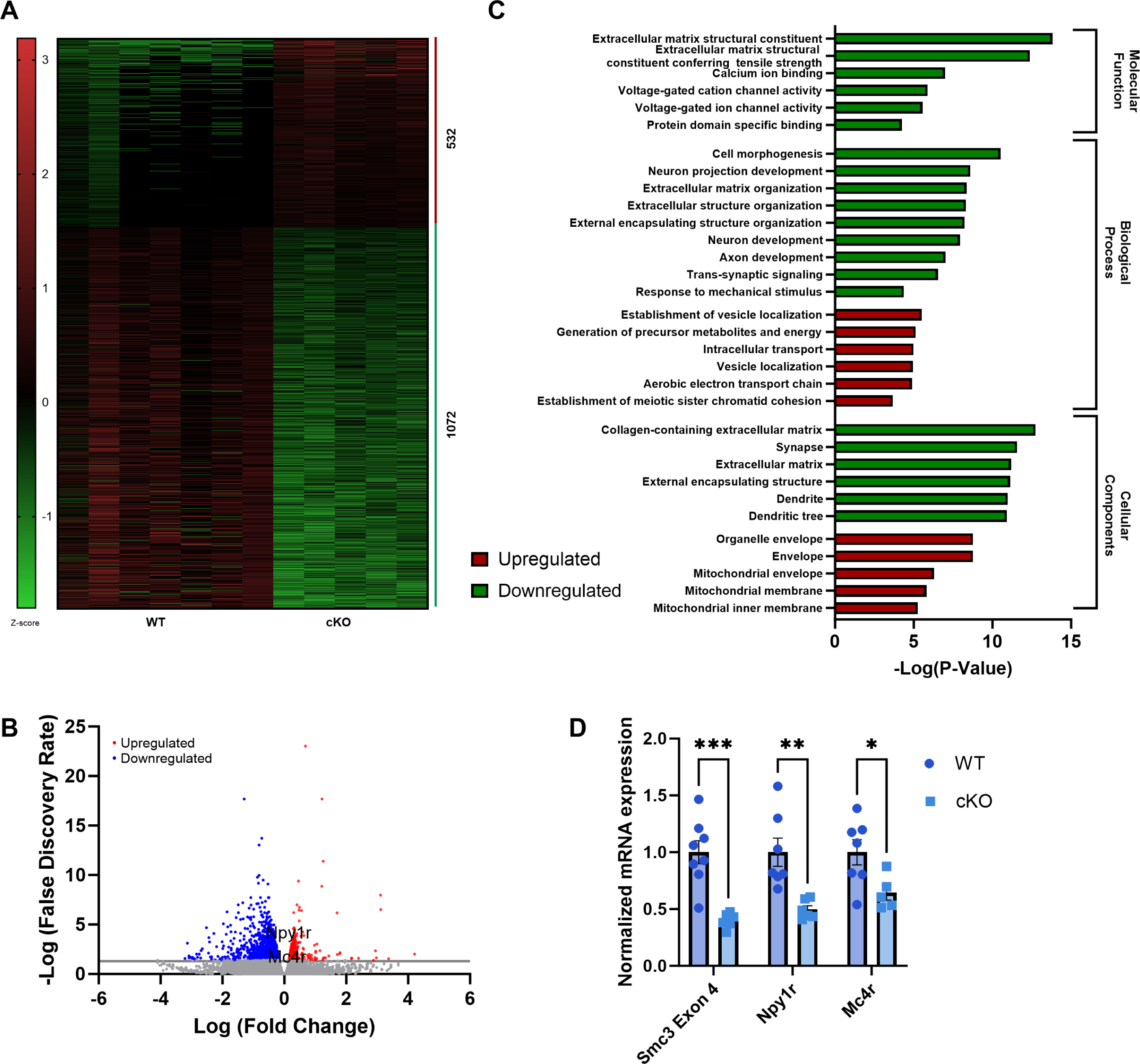
Transcriptomic analysis of hypothalamus in *Smc3* knockout mice. **(A)** Heatmap of differentially expressed genes in hypothalamus compares cKO mice to WT littermates 6 weeks after tamoxifen treatment— 532 upregulated genes and 1604 downregulated genes in *Smc3* knockout mice (FDR≤0.05; WT, n=7; cKO, n=5). **(B)** Volcano plot shows distribution of differently expressed genes in *Smc3* cKO mice. **(C)** Gene ontology analysis of differentially expressed genes in hypothalamus. **(D)** Quantitave Real-time PCR validation of RNA-seq data. (two-sided *t*-test WT, n=8; cKO, n=6; ***P<0.001, **P<0.01, *P<0.05)

Downregulated genes included *Mc4r* and *NPY1r*, two receptors highly involved in appetite and metabolic processes, and we verified these differences with real time PCR (Fig. 5D). Together, these data provide collective support for the morphological observations indicating dysregulation of dendritic structure in the hypothalamus and the corresponding observed phenotype.

Since SMC3 is a chromatin binding factor that can regulate gene transcription, we further set out to determine if gene expression changes may be due to changes in SMC3 binding in those genetic regions. In order to investigate genome-wide alterations in the binding dynamics of *Smc3* cKO mice (compared to wild-type), we conducted ChIP-seq assays on hypothalamic tissues. Our analysis unveiled 1681 sites significantly modified in the cKO group (Fig. 6A). We further categorized the results into three groups: sites that were ‘lost’ (no peak in cKO), ‘gained’ (no peak in WT), and sites that exhibited increased or decreased SMC3 -binding in cKO compared to WT. Consistent with our findings, a majority of the identified sites were either lost (604 sites) or exhibited decreased binding (517 sites) in the cKO mice (Supplementary Table 2).

**Figure 6.**
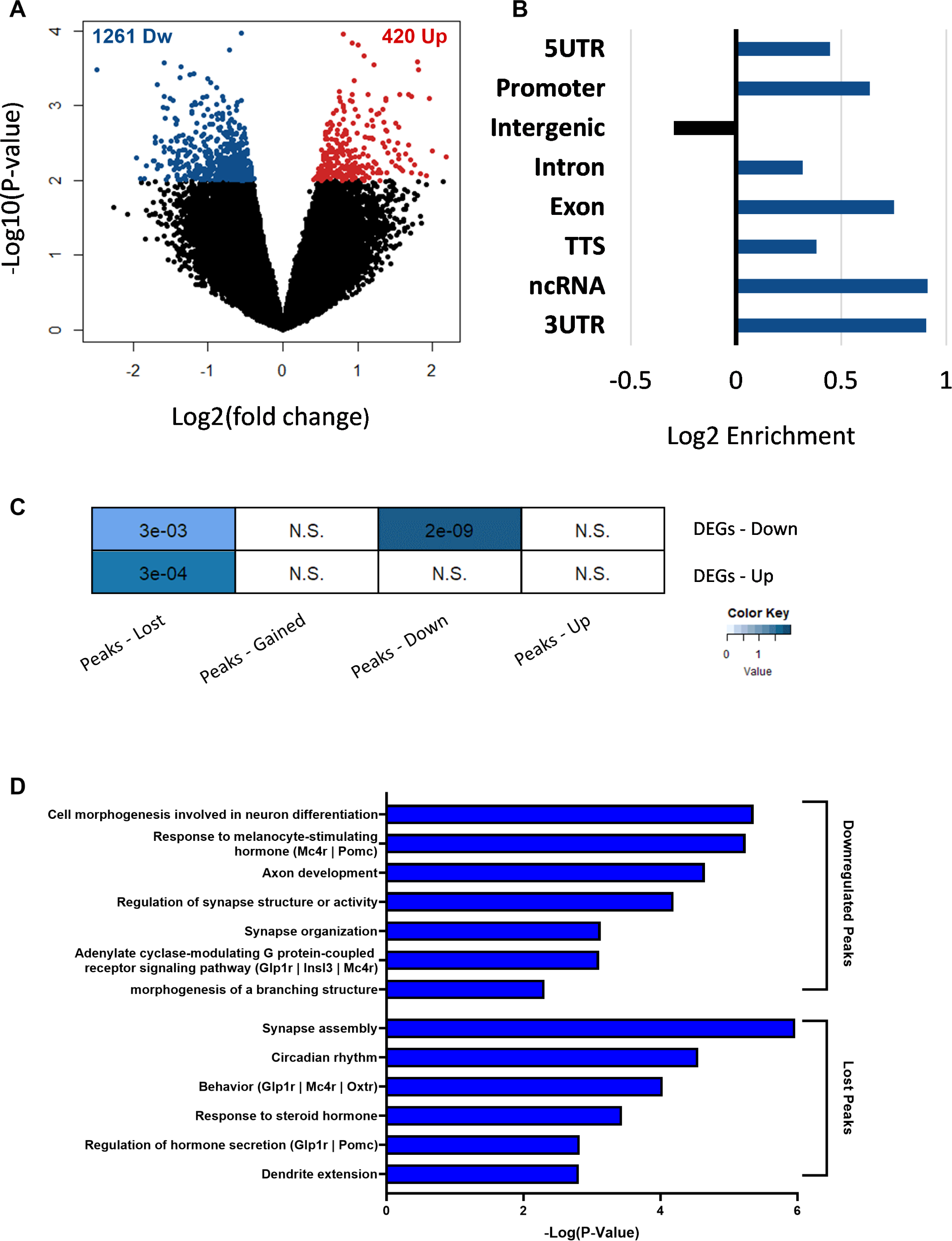
SMC3 ChIP-seq. reveals differential binding affinity in excitatory cell specific regions. **(A)** Differential binding affinity of SMC3 are presented in a volcano plot. Pairwise comparisons WT vs. cKO, P-value < 0.01; n = 5 biologically independent samples. **(B)** Annotation analysis (via HOMER tools) for all SMC3 differential bound sites. Numbers on the y-axis indicate the Log2 ratio values of observed loci (number of differential sites) over expected (total size (bp)) of the 5UTR, Promoter, Intergenic, Intron, Exon, TTS, ncRNA, 3UTR. **(C)** Correlation analysis was performed between DEGs and SMC3 -differential bound sites, which were parsed to gained, lost, down or upregulated peaks. P-value (presented in numbers) and odds ratio (color scale) from Fisher’s exact test were calculated by GeneOverlap package and presented in the heatmap. **(D)** Gene ontology (GO) analysis of gained, lost, down or upregulated peaks SMC3 - differential bound sites. GO analysis was performed using Metascape and values are presented in Log(P value) scale.

Differential SMC3 binding sites were enriched in genomic regions including promoters, exons and 3-UTR regions, and were underrepresented in intergenic regions (Fig. 6B).

Correlation analysis between differential Smc3 binding sites (identified through ChIP-seq) and differential genes (identified through RNA-seq) demonstrated a significant overlap (Fig 6C, GeneOverlap, Fisher’s exact test for P-value and odds ratio). Specifically, there was a pronounced significant correlation between lost/decreased binding sites and downregulated genes, which further underscores the pivotal role of SMC3 as a key regulator of gene expression in the hypothalamus (Fig 6C).

Gene ontology analysis of lost and decreased binding sites revealed many pathways consistent with our morphological and behavioral data, including axon development, several synaptic categories, circadian rhythms, and response to melanocyte-stimulating hormone. Decreased binding levels of SMC3 in regulatory regions assigned to the Mc4r gene were also found in these categories, suggesting that these genomic sites are specific to excitatory neurons and potentially influencing gene expression.

MC4 receptor in the hypothalamic Paraventricular nucleus (PVN) region is a known central regulator of food intake and metabolic processes^17,18^. Increases of peripheral hormones tested in Fig. 3 lead indirectly to activation of this receptor, and decreased appetite. Therefore, MC4r dysregulation mediated by Smc3, could be a putative pathway leading to the overweight phenotype. Immunohistochemistry verified a significant decrease of MC4r in the hypothalamus of *Smc3* cKO mice (Fig. 7).

**Figure 7.**
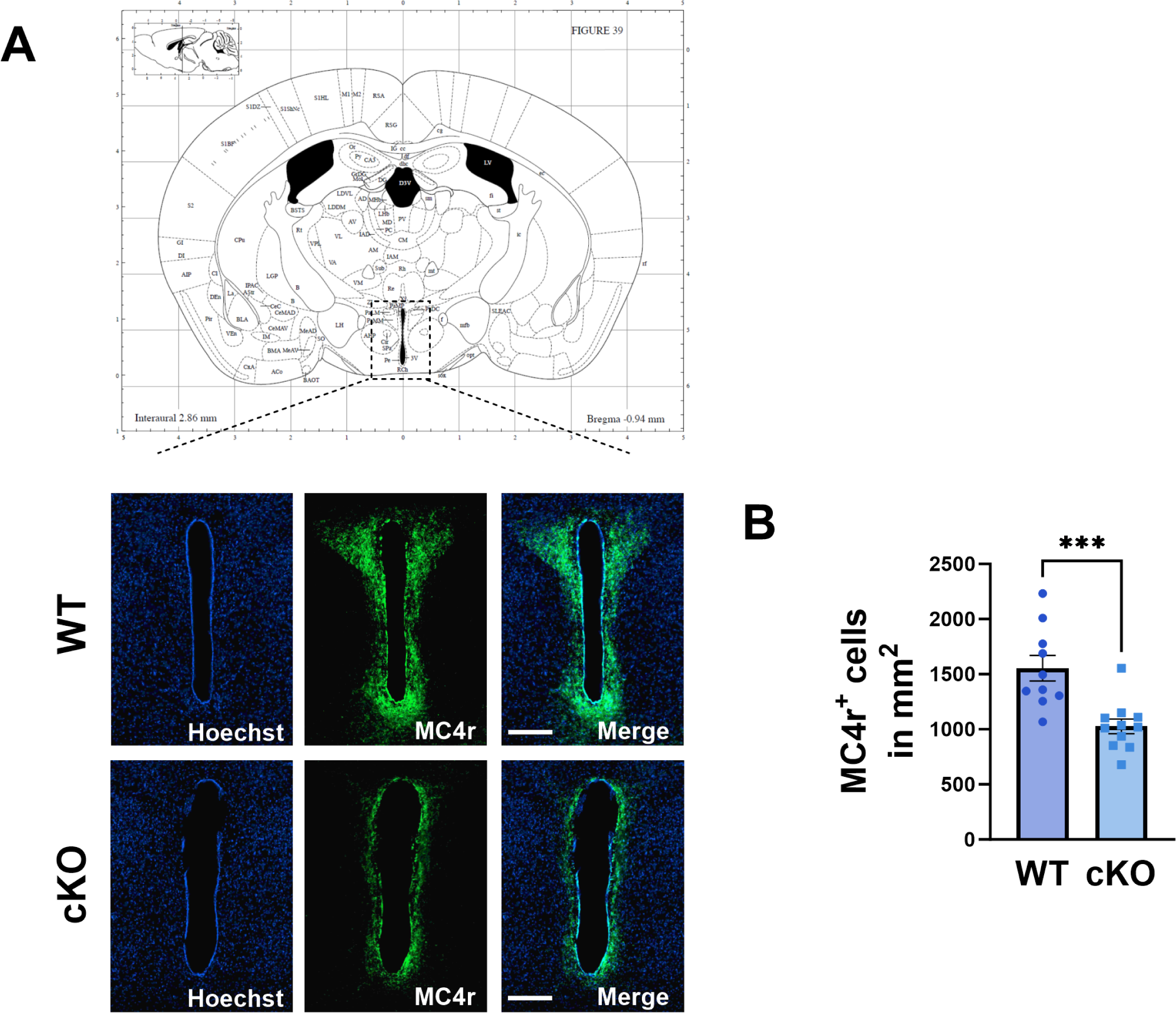
SMC3 regulates Melanocortin-4 receptor in hypothalamus. **(A)** Illustration adopted from mouse brain atlas^42^ indicate region of interest (ROI) that were stained for MC4 receptor. Representative hypothalamus of WT mouse and cKO mouse. Scale bars present 200*μm*. **(B)** Quantification of MC4r positive cells (two-sided *t*-test WT, n=10; cKO, n= 12; ***P=0.0007) in 1 mm^2^.

### Treatment of phenotype with Setmelanotide

To understand if MC4r signaling may be directly related to the overweight phenotype, we treated both wild type and *Smc3* cKO mice with Setmelanotide, a MC4r agonist. Setmelanotide was FDA approved for obesity disorder treatment by attenuating appetite^19^. Six weeks after tamoxifen treatment, wild type and *Smc3* cKO mice were treated with Setmelanotide I.P. (1 mg/kg/day), or vehicle control, for up to 12 days (Fig. 8A).

**Figure 8.**
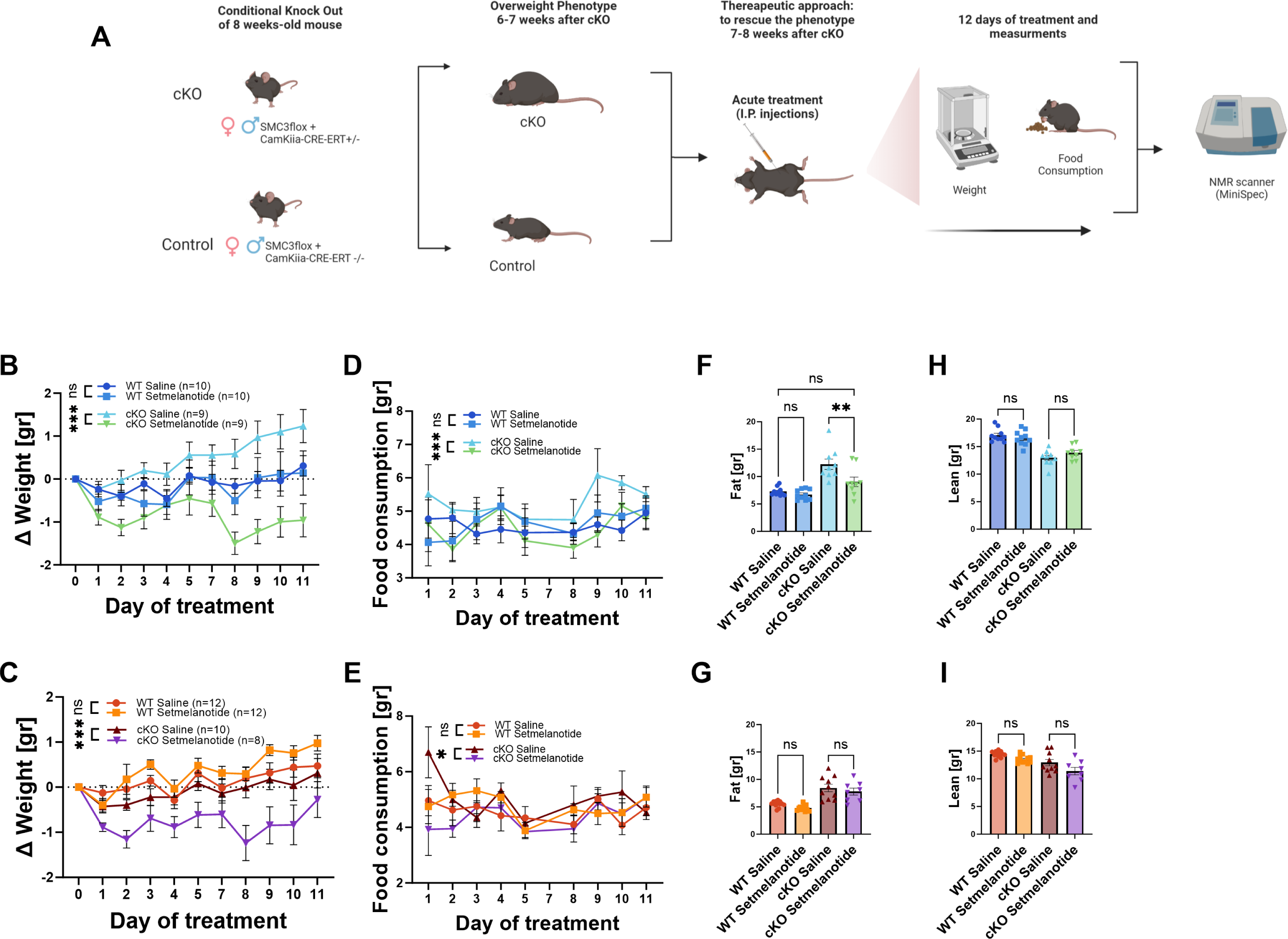
Setmelanotide treatment reverses metabolic phenotypes in *Smc3* cKO mice. **(A)** Schematic illustration **(B)** Weight measurements of male mice, presented as change of weight (Δweight) shows reduction of weight at cKO group that treated, statistically significant when compared to cKO that treated with Saline, moreover when compared to WT groups (One-Way ANOVA WT, n=10; cKO, n=9; F_3,30_=29.99; ***P<0.001, Tukey test). **(C)** Weight measurements of female mice, presented as change of weight (Δweight) shows reduction of weight at cKO group that treated, statistically significant when compared to cKO that treated with Saline, moreover when compared to WT groups (One-Way ANOVA WT, n=12; cKO - saline, n=10; cKO – Setmelanotide, n=8 F_3,30_=43.55; ***P<0.001, Tukey test). **(D)** Food consumption was measured individually for each mouse, the graph present daily intake. Significant difference was detected between two cKO groups, where the treated with Setmelanotide mice shows reduce in food consumption (One-Way ANOVA WT, n=10; cKO, n=9; F_3,24_=10.34; ***P<0.001, Tukey test). **(E)** Food consumption was measured individually for each mouse, the graph present daily intake. Significant difference was detected between two cKO groups, where the treated with Setmelanotide mice shows reduce in food consumption (One-Way ANOVA WT, n=12; cKO - saline, n=10; cKO – Setmelanotide, n=8; F_3,24_=2.855; *P<0.05, Tukey test). **(F)** Body composition of Fat tissue, that measured using Mini-spec system; at last day of treatment (day 12). The result show that cKO mice decreased their fat tissue after Setmelanotide treatment, with no significant difference between WT littermates. (One-Way ANOVA WT, n=10; cKO, n=10; F_3,34_=14.84; ns P>0.05 not significant; **P=0.008, Tukey test). **(G)** Body composition of Fat tissue, that measured using Mini-spec system; at last day of treatment (day 12). The result show no significant difference between all four groups. (One-Way ANOVA WT, n=12; cKO - saline, n=10; cKO – Setmelanotide, n=8; F_3,27_=18.00; ns P>0.05 not significant, Tukey test). **(H)** Body composition of lean tissue, measured at the same as fat; shows the same pattern of decreased lean tissue at cKO group, and without any significant changes due to Setmelanotide treatment (One-Way ANOVA WT, n=10; cKO, n=10; F_3,34_=22.41; ns P>0.05 not significant, Tukey test). **(I)** Body composition of lean tissue, measured at the same as fat; shows absence of any significant changes due to Setmelanotide treatment (One-Way ANOVA WT, n=12; cKO - saline, n=10; cKO – Setmelanotide, n=8; F_3,38_=9.449; ns P>0.05 not significant, Tukey test).

*Smc3 cKO* mice treated with Setmelanotide displayed significantly decreased weight gain compared to all other experimental groups in both sexes (Fig. 8B, C). Food consumption was decreased in the knockout-Setmelanotide group compared to the knockout-Saline group in both sexes (Fig. 8D, E). In male mice, fat tissue was also increased in the Saline-*Smc3* cKO group compared to the wild type groups, and Setmelanotide-*Smc3* cKO group displayed lower fat than the Smc3-cKO group (Fig. 8F). Females did not show any change in the fat tissue (Fig. 8G). Lean tissue was not affected in any of the experimental groups in males or females (Fig. 8H, I). Therefore, Setmelanotide was able to attenuate many of the metabolic changes in the *Smc3* cKO animals.

## Discussion

Most previous studies have found that depletion of *Smc3* is necessary for cell survival and proper development^6^. Fujita et. al. demonstrated that neuron-specific homozygous knockout of *Smc3* during development induced postnatal fatality. In our study, where *Smc3* was removed during adulthood, mice survived with no detectable apoptosis. Therefore, *Smc3* knockout in development leads to neuronal death, while knockout in postmitotic adulthood neurons does not. This highlights that *Smc3* plays different roles during development and adulthood.

CdLs is often associated with undergrowth during development until teenage years^8,20^. Some studies, including a large study of individuals with CdLs in Italy, have shown that individuals become overweight during adulthood. Fujita et. al. demonstrated that neuron-specific *Smc3* haploinsufficient mice do not display any significant weight changes in the first eight weeks of development^11^. In our study, we find an increase in weight starting five weeks after *Smc3* depletion in the adulthood. There are a few possible explanations of differences between the two studies. First, it is possible that *Smc3* has an appetite suppressing effect specifically in adulthood. This explanation would correlate well with the human condition, where individuals with CdLs often show weight gain specifically in the adulthood. However, since we only determined the phenotype of *Smc3* homozygote knockout mice, it is possible that we would not see the same phenotype in a happloinsufficient model. One major limitation of the current study is that we have now yet determined the phenotype of adulthood *Smc3* haploinsufficiency.

*Smc3* conditional knockout mice displayed a robust increase in several appetite-suppressing hormones, including leptin, and GLP-1. This shows a normal peripheral attempt to suppress both appetite and weight gain. Leptin and GLP-1 both exert their effects on appetite regulation by acting on the hypothalamus. The hypothalamus is densely populated with Pro-opiomelanocortin (POMC) neurons, which express receptors and are sensitive to both hormones. The interaction between leptin and GLP-1 in the brain involves intricate signaling pathways that ultimately suppress appetite behavior and affect metabolism^21,22^. Therefore, we hypothesized that the hypothalamus may not be functioning properly and may not be responding to these signals. In fact, we found a reduction in dendritic arbor in hypothalamus as result of *Smc3* knockout.

Dendritic arborization is essential for receiving and integrating synaptic inputs from other neurons facilitating the transmission of electrical impulses and chemical signals across neuronal networks^23^. Therefore, decreased dendritic arborization supports the theory that the hypothalamic dysfunction may be involved in the metabolic phenotype. The finding of decreased dendritic arborization diverges from a previous study, which reported an increase of dendritic arborization in the cortex as result of haploinsufficient *Smc3* knockout at the embryonic stage^11^. These differences highlight how the timing of gene knockout and the specific brain area being affected by neuronal-specific lack of SMC3leads to different phenotypes at the cellular level.

An analysis of differentially expressed genes in *Smc3* conditional knockout mice revealed significant downregulation of genes involved in biologically important processes for neuronal maintenance, such as dendritic trees, synapses, axons, neuron development, and channel activity. These findings correlate well with the morphological changes of the dendritic arboring. Surprisingly, there was an increase in expression of genes involved in mitochondrial function and oxidative phosphorylation. Dysregulatoin of mitochondrial function is been reported as part neurodegenerative diseases like Alzheimer and Parkinson^24–26^. Efficient mitochondrial energy production, maintenance of calcium homeostasis, protection against oxidative stress, and involvement in synaptic function collectively contribute to optimal neuronal health and brain function^27^. Of particular interest, hyperactive mitochondria are part of the pathology of Parkinson’s Disease, and therapeutic options are being to developed to reverse the effects of hyperactive neuronal mitochondria^28^.

Previous studies of cohesin subunits knockout in neurons were performed in cell culture. *Calderon et. al.* determined that lack of cohesin subunit Rad21 in immature postmitotic neurons induced downregulation in 1028 genes and upregulation in 572 genes^29^, which correlates strongly with our findings of knockout of neuronal Smc3 in the adulthood (1072 downregulated and 532 upregulated). Among the downregulated genes, both studies also revealed very similar gene ontology categories, relating to synapses, neuronal structure, and ion channels or transport. Therefore, both studies verify that the cohesin complex has a very robust role in gene expression in postmitotic neurons, and further suggest that the complex has a larger role in promoting gene expression.

The ChIP-seq results from our study provide additional evidence supporting the notion that Smc3 plays a substantial role in the neuronal genome, contributing to the maintenance of gene expression. For example, Smc3 binding sites that were downregulated or lost in cKO mice, were predominantly associated with excitatory neuron signatures and overlapped with genes that were downregulated in the Smc3 cKO mice. Further studies would need to be constructed to consider if excitatory neuron-specific Smc3 binding changes the three-dimensional chromatin structure in neurons, and to determine the mechanism of how this may maintain gene expression.

Among downregulated genes, we identified Melanocortin 4 receptors (MC4r), a receptor heavily expressed in the hypothalamus^30^. Activation of MC4r by its endogenous ligands, melanocortins, suppresses appetite and promotes energy expenditure, contributing to the maintenance of body weight homeostasis^31–33^. Mutations in the MC4r gene have been identified as a leading cause of monogenic obesity in humans^34^ while activating this receptor reported as efficient way to fight obesity^35^.

Setmelanotide functions as a selective MC4r agonist, directly binding to and activating the MC4r located within the hypothalamus. By mimicking the natural ligands that activate MC4r, Setmelanotide enhances the signaling cascade responsible for appetite regulation and energy expenditure. Setmelanotide was recently granted approval by the U.S. Food and Drug Administration (FDA)^19^ Efficacy of Setmelanotide was determined in individuals with rare genetic mutations that disrupt the MC4r pathway such as POMC^36^. Of interest, a recent study found that Setmelanotide can also affect treat individuals with dysregulated methylation in the POMC gene, which further suggests that epigenetic dysregulation of this pathway may be treated with Setmelanotide^37^. A limitation of the current study is that we do not know which specific cell types in the hypothalamus are responsible for the phenotype. Several cell types in the hypothalamus integrate signals and work in tandem to regulate appetite, and it is not currently clear which of these cell types is responsible for the observed metabolic phenotype.

Through the administration of Setmelanotide, we achieved reduction in body weight and food consumption among both male and female mice. Interestingly, Setmelanotide had effects specifically on the *Smc3* conditional knockout mice, without having effects on wild type mice. A previous study found that Magel2 knockout mice, which have deficiencies in the POMC-expressing neurons, were also particularly sensitive to Setmelanotide treatment^38^. This suggests that under certain conditions, animals that are deficient in the POMC/Mcr4 pathway can be treated with Setmelanotide. To conclude we determined that SMC3 regulates gene expression in the adult hypothalamus and adulthood neuronal SMC3 is an important regulator of metabolic processes.

## Materials and Methods

### Mouse model

All experimental procedures were conducted in accordance with the Federation of Laboratory Animal Science Associations (FELSA) and the National Institutes of Health regulations and were approved by the Bar-Ilan University Institutional Animal Care and Use Committee (IACUC), protocols ID: BIU-MD-IL-2206-151-3 and BIU-MD-IL-2301-102-5.

The mice were bred and housed in a vivarium in Plexiglas cages, with constant temperature and humidity (22°C, 50%) 12-hour light\dark cycle, food (#1234 TPF, Altromin, Lage, Germany) and water provided ad libitum.

To generate post-mitotic neuronal-specific *Smc3* knockout mice, C57/Bl6 *Smc3* loxp/loxp (floxed *Smc3*) female mice with were crossed with C57/Bl6 males that are floxed *Smc3* and Camk2a-cre/ER^T^2^+^ (Cre+). Therefore, offspring are all floxed *Smc3*, and half are Camk2a-cre/ER^T^2^+^ (Cre^+^) while half are Camk2a-cre/ER^T^2^-^ (Cre^-^). Mice with Cre^+^ named as cKO due the lack of *Smc3* gene after tamoxifen treatment. Mice with Cre^-^ served as WT. All mice were treated with tamoxifen. The CAMKIIa-Cre-ERT line was kindly provided by Prof. Alon Chen (Weizmann Institute of Science)^13^. Floxed *Smc3* animals were purchased from Jackson Laboratory (stock: 030559) (Bar Harbor, Maine, USA)^39^.

In order to reduce and refine our animal research, metabolic chambers, hormonal analysis and RNA-seq were tested only on male mice.

### Immunohistochemistry

Mice were injected with veterinary Pentobarbital sodium – Pental 160 mg/kg (CTS Chemical Industries, Ltd, Hod-Hasharon, Israel, #077299195200), perfused with 4% paraformaldehyde followed by perfusion with saline. Brains were dissected and fixed overnight at 4°C in 4% paraformaldehyde, then incubated with 30% sucrose. Brains were sectioned to 30*μm* floating slices by sliding microtome. Slices were blocked for one hour in blocking solution (10% horse serum, 10% triton and TBS). Incubation with primary antibodies performed overnight at room temperature. αSMC3 (Bethyl, Montgomery, Texas, USA, #A300-060A) (1:1000); NeuN (Merck Millipore, Darmstadt, Germany, #MAB377) (1:200); QKI (UC Davis/NIH NeuroMab Facility, #N1476/6) (1:200); Mc4r (Alomone labs, Jerusalem, Israel, #AMR-024) (1:200). Incubation for one hour at room temperature was made with secondary antibody: Goat αRabbit Alexa flour 488 (Invitrogen Rhenium, Modi’in, Israel, #A-11008) (1:200) or Goat αMouse Cy-3 (Invitrogen, #A10521) (1:500). At the end Hoechst (Sigma-Aldrich, Rehovot, Israel, #94403) (1:1,000) was used for 10 minutes, before covering with Shandon-mount (Thermo scientific, Rhenium, Modi’in, Israel, #1900331). Immunohistochemistry images were taken with Zeiss AxioImager M2 with the apotome system or Axio Scan.Z1 microscope (Zeiss, Jena, Germany), processed with Zen pro software (version 3.3.89.0000, windows).

### Tunnel staining

After 4 weeks of tamoxifen treatment, 30 *μm* sections were stained according to instruction of *In situ* Cell Death Detection Kit (Roche, Basel, Switzerland, #11684817910). Briefly, brain slices were permeabilized by 0.3% triton in PBS and positive control slices were incubated with Dnase I recombinant (Roche, Basel, Switzerland, #4536282001) (1,000 units). Labeled slices were stained in the end with Hoechst solution (Sigma-Aldrich, Rehovot, Israel, #94403) (1:1,000) for 10 minutes and then mounted on slides (Thermo scientific, Rhenium, Modi’in, Israel, Shandon-mount #1900331). Images were taken with Zeiss AxioImager M2 with the apotome system (Zeiss, Jena, Germany).

### Golgi staining and sholl analysis

Fresh dissected brains were stained according to instruction of SuperGolgi kit (Bioenno Tech, LLC, Santa Ana, California, USA, #003010). Briefly, brains were washed with PBS and placed for two weeks of impregnation. Coronal sections of the brains to 175*μm* was performed using vibrotome Leica VT 1200s. Sections were mounted to glass slides and staining using the kit reagents following manufacturer’s instructions.

Stack images of neurons from hypothalamus and CA1 region of hippocampus were acquired using Zeiss AxioImager M2 (Zeiss, Jena, Germany) with apotome system (with ORCA-Flash 4.0 V3 camera). Images were acquired as Z-stack with 0.5 *μm* intervals at 40× magnification.

Dendritic arborization (sholl analysis) calculated via automated filament reconstruction in Imaris (version 9.9 windows).

### RNA-seq and Gene ontology

RNA from frozen brain punches was purified using the Rneasy mini kit (Quigen, Ilex, Petah-Tikva, Israel, # 74004) according to the manufacturer’s instructions. Quality of isolated RNA was tested using the Agilent RNA Pico 6000 kit and Bioanalyzer 2100 at the Genome Technology Center of the Azrieli Faculty of Medicine, Bar-Ilan University. 350 ng of total RNA were used for mRNA enrichment using NEBNext mRNA polyA isolation module (NEB, Ornat, Rehovot, Israel, #E7490L) and libraries for Illumina sequencing were prepared using the NEBNext Ultra II RNA kit (NEB, Ornat, Rehovot, Israel, #E7770L). Quantification of the library was performed using dsDNA HS assay kit and Qubit (Molecular Probes, Life Technologies, Rhenium, Modi’in, Israel) and quantification was performed using the Agilent DNA HS kit and Bioanalyzer 2100. 10 nM of each library were pooled together and were diluted to 4 nM according to NextSeq manufacturer’s instructions. 1.1 pM was loaded into the flow cell with 1% PhiX library control. Reads were mapped to the Mus musculus reference genome (mm9) using the Tophat2 software (release Tophat2.0.12). Differential gene expression analysis was performed using the DESeq2 pipeline.

Gene ontology analysis on differentially expressed genes was performed using the Toppgenetool (https://toppgene.cchmc.org/) ^38^ .

Accession numbers for the raw data files for RNA-seq analysis reported in this paper is GEO: GSE243287.

### Metabolic chambers

Respiratory exchange rate (RER), heat production and distance were measured and calculated by PhenoMaster caging system (TSE system, Labotal, Abu Gosh, Israel, TSE5403). Mice were put in a single cage of the TSE system, after three days of habituation, measurements were taken every 40 minutes, under 12-hour light\dark cycle, food and water provided ad libitum.

### Body composition scanning

Body composition (fat and lean) was measured using pulsed time-domain nuclear magnetic resonance (NMR; model LF50, Bruker, Rehovot, Israel). Unanesthetized mice were placed in a restrainer tube that is inserted into the system for approximately 2 min. The mouse was then returned to its home cage.

### Quantitative real-time PCR (qRT-PCR)

RNA extracted from WT and cKO mice hypothalamus, using Rneasy Mini kit (Qiagen, #74104). RNA was quantified using NanoDrop TM 2000 (Thermo Scientific, Rhenium, Modi’in, Israel). Two micrograms of total RNA were reverse transcribed using High-Capacity RNA to cDNA kit (Applied Biosystems, Rhenium, Modi’in, Israel, #4368814). Quantitative PCR analysis was performed using or FAST SYBR Green master mix (Applied Biosystems, Rhenium, Modi’in, Israel, # AB-4385612) in triplicates using the ViiA7 real-time PCR system (Applied Biosystems, Rhenium, Modi’in, Israel). PCR program was 95°C for 10 min, with forty cycles of denaturation at 95°C for 10 sec, and annealing/elongation at 60°C for 30 sec. Results were quantified using the comparative Ct (ΔΔCt) method for Hprt gene. Primer specificity was verified by running qPCR product in 3% agarose gel and visualization of appropriate band.

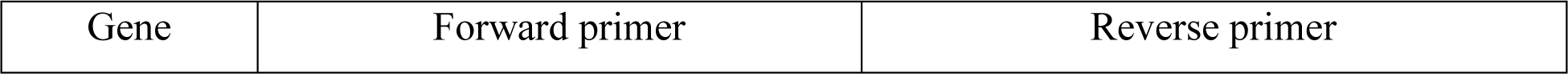
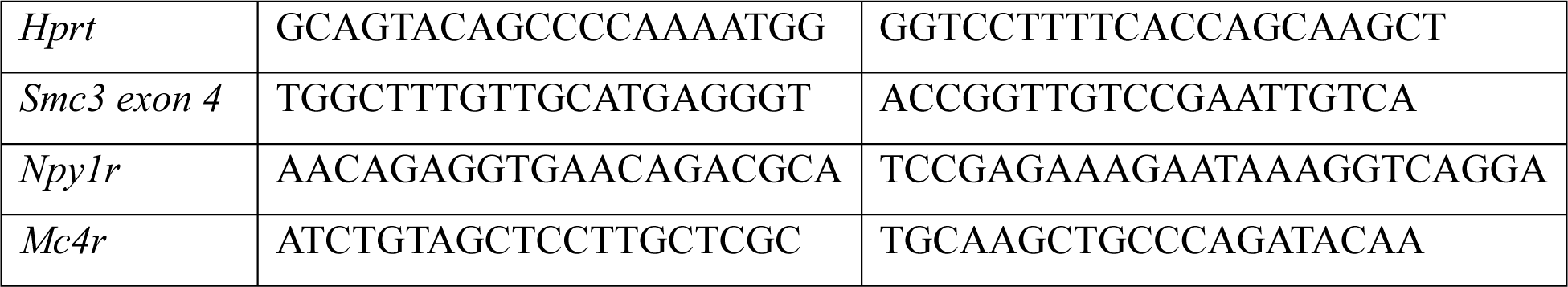

### Setmelanotide treatment

Setmelanotide (BOC Sciences, Shirley, NY, USA, #920014728) was used for treatment. When mice reached the appropriate weight, treatment performed by intraperitoneal injection once a day for 12 days with Setmelanotide (1 mg/kg) or saline^40,41^.

### Hormonal analysis

Hormonal analysis was performed using MILLIPLEX mouse metabolic hormone expended panel (Merck Millipore, Darmstadt, Germany, #MMHE-44K). Up to 0.2ml of blood was collected from mouse tail vein. After centrifugation, plasma extract was moved into sterile tubes that contained 0.5M EDTA, DPP-IV (Sigma-Aldrich, Rehovot, Israel, #PHR1857-1G), Pefabloc (Roche, Basel, Switzerland, #30827997) and Halt Protease and phosphatase inhibitor (Thermo scientific, Rhenium, Modi’in, Israel, #778442). The staining of the samples was performed according to the manufacturer’s instructions and loaded into MAGPIX system (Luminex corporation, Austin, Texas, USA). Data were analyzed using column statistic in GraphPad Prism (Windows, version 9.5).

### Chromatin immunoprecipitation (ChIP) sequencing

To comprehensively identify both the binding sites and enrichment levels of SMC3 across the genome, we employed chromatin immunoprecipitation (ChIP) followed by sequencing assay on the hypothalamic tissue from WT and cKO mice. ChIP-seq was performed independently on five wild type and five *Smc3* cKO hypothalami. Briefly, each cross-linked hypothalamic sample (1% formaldehyde) was sonicated in cell lysis buffer, resulting in chromatin fragments ranging from 200 to 1000 bp. These fragments incubated in ChIP dilution buffer with an αSMC3 antibody (5 μg/sample; Bethyl, Montgomery, Texas, USA, #A300-060A). Simultaneously, 5% input (without immunoprecipitation step) was utilized to normalize the amount of chromatin in each sample. Subsequently, DNA was isolated from each immunoprecipitated/input sample and subjected to genomic library preparation at the Azrieli Faculty of Medicine Genome Center (Bar Ilan University). 20*μL* of DNA were used for libraries preperation using the NEBNext Ultra II DNA kit (NEB, #E7645). Quantification of the library was performed using dsDNA HS Assay Kit and Qubit 2.0 (Molecular Probes, Life Technologies) and qualification was done using the Agilent D1000 Tapestation Kit and Tapestation 4200. 25nM of each library was pooled together and was diluted to 4nM according to NextSeq manufacturer’s instructions. 1.5pM was loaded onto the Flow Cell with 1% PhiX library control. Libraries were sequenced on an Illumina NextSeq 500 instrument, 75 cycles single read sequencing. Following bioanalyzer - QC for library size and distribution, the libraries will be sequenced on the Illumina Nextseq 550 platform. Raw sequencing data from Chip-seq analysis is available at GSE252366.

ChIP-seq analysis included the aligning of the raw fastq reads to the mouse mm9 reference genome using Bowtie 2.0. The Samtools collection used to sort, remove PCR duplicates (rmdup), index BAM files (index), and calculate library statistics. SMC3 ‘peaks’ determined by the MACS2 package (callpeak function, --broad peaks). Lastly, differential binding analysis was performed where immunoprecipitated over input scores were calculated using the Diffbind package (cutoff P-value<0.01). Differential peaks were divided into the following categories according to their binding affinity score; (i) lost peaks (score in WT – 0 in cKO), (ii) gained peaks (0 in WT – score in cKO), (iii) up and (iv) downregulated peaks. Correlation analysis between differential SMC3 binding sites and DEGs was performed by GeneOverlap R package.

### Statistical analysis

Data were analyzed with GraphPad Prism (Windows, version 9.5). *In-vivo* data and Golgi staining for morphology are presented as mean ± standard error of the mean (SEM), when *n* values present biological repeats. Different statistical tests were used as appropriate for each experiment. The statistical significance was calculated using unpaired two-tailed *t*-test with confidence level of 95%. One-way ANOVA and Two-way ANOVA followed by Tukey-Kramer or Dunnett’s posthoc test. Third-order polynomial Non-linear regression curve was performed for dendritic branching of sholl analysis, with confidence level of 99%.

All values were considered as significant if P<0.05 — For all the figures *P<0.05, **P<0.01, ***P<0.001, ****P<0.0001, ns P>0.05 not significant.

## Acknowledgements

We thank the Azrieli Faculty of Medicine Genome Center (Bar Ilan University) for help with sequencing. We thank Dr. Aviad Sivan for help with bioinformatic analysis in the RNA-seq analysis and Dr. Gilgi Friedlander from the Israel National Center for Personalized Medicine in the Weizmann Institute of Science. In addition, we thank Dr. Roey Lahav and the team at the Azrieli Faculty of Medicine Preclinical animal facility for assistance with metabolic cages and general animal care. This study was funded by grants from the Israel Science Foundation (898/17 and 1159/22).

## Author Contributions

N.S. performed the experimentation and analysis outlined in this manuscript and wrote the manuscript. D.G. performed behavioral analysis. J.B. performed Chip-seq experimentation. R.G. analyzed metabolic data. A.M. performed analysis on Chip-seq and RNA-seq datasets. E.E. supervised the project, obtained funding and wrote the manuscript. All authors edited the manuscript.

## Declaration of Interests

The authors declare no competing interests.

## Supplementary material

**Supplementary Figure 1. Change in body composition expressed before the weight gain**

As result of obesity phenotype, mice were measures at Mini-spec system for body composition of fat and lean tissue at day 28 after cKO induced. **(A–B)** Shows the measurements of male mice, at day 28. **(C–D)** Shows the measurements of female mice. Both, males and females, shows the induce in fad body at the cKO groups, and reduce of lean tissue. That can be observed even before the obesity is significant between WT and cKO groups in weight measurements.

**Supplementary Figure 2. Hormonal changes at 9 weeks after cKO of *Smc3* that leading to dysregulated behavioral**

As part of understanding the late phenotype of *Smc3* cKO, we measured the mice not only at 6 weeks after cKO induced (presented at Figure 3) but also at 9 weeks. A-F presents the same pattern of dysregulation appetite hormones as it measured at 6 weeks, with the addition of Glucagon measurements (F) that calculated as significant (two-sided *t*-test; n=9; *P<0.05, **P<0.01, ***P<0.001, ****P<0.0001, ns P>0.05 not significant).

**Supplementary Figure 3. Knockout of *Smc3* gene in excitatory neurons of hypothalamus**

(**A)** Representative pictures of hippocampus in wild type and *Smc3* knockout mice 5 weeks after tamoxifen treatment.; Scale bars present 500*μm*. **(B)** quantification of SMC3 positive cells in 1 mm^2^ hypothalamus both groups (two-sided *t*-test WT, n=4; cKO, n=5; *P=0.0228).

## References

1. Nasmyth, K. & Haering, C. H. Cohesin: Its Roles and Mechanisms. Annu. Rev. Genet. 43, 525–558 (2009).

2. Wong, R. W. An update on cohesin function as a ‘molecular glue’ on chromosomes and spindles. Cell Cycle 9, 1754–1758 (2010).

3. Piché, J., Van Vliet, P. P., Pucéat, M. & Andelfinger, G. The expanding phenotypes of cohesinopathies: one ring to rule them all! Cell Cycle 18, 2828 (2019).

4. Liu, J. & Krantz, I. D. Cornelia de Lange syndrome, cohesin, and beyond. Clin. Genet. 76, 303–314 (2009).

5. Gil-Rodríguez, M. C. et al. De Novo Heterozygous Mutations in SMC3 Cause a Range of Cornelia de Lange Syndrome-Overlapping Phenotypes. Hum. Mutat. 36, 454–462 (2015).

6. Kline, A. D. et al. Diagnosis and management of Cornelia de Lange syndrome: first international consensus statement. Nat. Rev. Genet. 2018 1910 19, 649–666 (2018).

7. Jackson, L., Kline, A. D., Barr, M. A. & Koch, S. de Lange syndrome: a clinical review of 310 individuals. Am. J. Med. Genet. 47, 940–946 (1993).

8. Mariani, M. et al. Adolescents and adults affected by Cornelia de Lange syndrome: A report of 73 Italian patients. Am. J. Med. Genet. Part C Semin. Med. Genet. 172, 206–213 (2016).

9. White, J. K. et al. Genome-wide Generation and Systematic Phenotyping of Knockout Mice Reveals New Roles for Many Genes. Cell 154, 452 (2013).

10. Fujita, Y. & Yamashita, T. Spatial organization of genome architecture in neuronal development and disease. Neurochem. Int. 119, 49–56 (2018).

11. Fujita, Y. et al. Decreased cohesin in the brain leads to defective synapse development and anxiety-related behavior. J. Exp. Med. 214, 1431–1452 (2017).

12. Weiss, F. D. et al. Neuronal genes deregulated in Cornelia de Lange Syndrome respond to removal and re-expression of cohesin. Nat. Commun. 12, (2021).

13. Erdmann, G., Schütz, G. & Berger, S. Inducible gene inactivation in neurons of the adult mouse forebrain. BMC Neurosci. 8, (2007).

14. Zha, C., Gamache, K., Hardt, O. M. & Sossin, W. S. Behavioral characterization of Capn15 conditional knockout mice. Behav. Brain Res. 454, 114635 (2023).

15. Kim, M. S. et al. An Essential Role for Histone Deacetylase 4 in Synaptic Plasticity and Memory Formation. J. Neurosci. 32, 10879–10886 (2012).

16. Lipinski, M. et al. KAT3-dependent acetylation of cell type-specific genes maintains neuronal identity in the adult mouse brain. Nat. Commun. 11, (2020).

17. Hinney, A., Volckmar, A. L. & Knoll, N. Melanocortin-4 receptor in energy homeostasis and obesity pathogenesis. Prog. Mol. Biol. Transl. Sci. 114, 147–191 (2013).

18. Balthasar, N. et al. Divergence of melanocortin pathways in the control of food intake and energy expenditure. Cell 123, 493–505 (2005).

19. Hussain, A. & Farzam, K. Setmelanotide. StatPearls (2023).

20. Kline, A. D. et al. Natural history of aging in Cornelia de Lange syndrome. Am. J. Med. Genet. C. Semin. Med. Genet. 145C, 248–260 (2007).

21. Kabahizi, A. et al. Glucagon-like peptide-1 (GLP-1) signalling in the brain: From neural circuits and metabolism to therapeutics. Br. J. Pharmacol. 179, 600–624 (2022).

22. Morrison, C. D. Leptin signaling in brain: A link between nutrition and cognition? Biochim. Biophys. Acta 1792, 401–408 (2009).

23. Jan, Y. N. & Jan, L. Y. Branching out: mechanisms of dendritic arborization. Nat. Rev. Neurosci. 11, 316–328 (2010).

24. Popov, L. D. Mitochondrial biogenesis: An update. J. Cell. Mol. Med. 24, 4892–4899 (2020).

25. Moradi Vastegani, S., et al. Mitochondrial Dysfunction and Parkinson’s Disease: Pathogenesis and Therapeutic Strategies. Neurochem. Res. 48, 2285–2308 (2023).

26. Jornayvaz, F. R. & Shulman, G. I. Regulation of mitochondrial biogenesis. Essays Biochem. 47, 69–84 (2010).

27. Rangaraju, V. et al. Pleiotropic Mitochondria: The Influence of Mitochondria on Neuronal Development and Disease. J. Neurosci. 39, 8200–8208 (2019).

28. Mor, D. E. et al. Metformin rescues Parkinson’s disease phenotypes caused by hyperactive mitochondria. Proc. Natl. Acad. Sci. U. S. A. 117, 26438–26447 (2020).

29. Calderon, L. et al. Cohesin-dependence of neuronal gene expression relates to chromatin loop length. Elife 11, 76539 (2022).

30. Wang, K. et al. Neuroanatomy of melanocortin-4 receptor pathway in the mouse brain. Open life Sci. 15, 580–587 (2020).

31. Fan, W., Boston, B. A., Kesterson, R. A., Hruby, V. J. & Cone, R. D. Role of melanocortinergic neurons in feeding and the agouti obesity syndrome. Nature 385, 165– 168 (1997).

32. Marie, L. S., Miura, G. I., Marsh, D. J., Yagaloff, K. & Palmiter, R. D. A metabolic defect promotes obesity in mice lacking melanocortin-4 receptors. Proc. Natl. Acad. Sci. U. S. A. 97, 12339–12344 (2000).

33. Namjou, B. et al. Evaluation of the MC4R gene across eMERGE network identifies many unreported obesity-associated variants. Int. J. Obes. (Lond). 45, 155–169 (2021).

34. Farooqi, I. S. & O’Rahilly, S. Monogenic obesity in humans. Annu. Rev. Med. 56, 443– 458 (2005).

35. Lotta, L. A. et al. Human Gain-of-Function MC4R Variants Show Signaling Bias and Protect against Obesity. Cell 177, 597–607.e9 (2019).

36. Trapp, C. M. & Censani, M. Setmelanotide: a promising advancement for pediatric patients with rare forms of genetic obesity. Curr. Opin. Endocrinol. Diabetes. Obes. 30, 136–140 (2023).

37. Lechner, L. et al. Early-set POMC methylation variability is accompanied by increased risk for obesity and is addressable by MC4R agonist treatment. Sci. Transl. Med. 15, (2023).

38. Chen, J., Bardes, E. E., Aronow, B. J. & Jegga, A. G. ToppGene Suite for gene list enrichment analysis and candidate gene prioritization. Nucleic Acids Res. 37, W305– W311 (2009).

39. Viny, A. D. et al. Dose-dependent role of the cohesin complex in normal and malignant hematopoiesis. J. Exp. Med. 212, 1819–1832 (2015).

40. Bischof, J. M., Van Der Ploeg, L. H. T., Colmers, W. F. & Wevrick, R. Magel2-null mice are hyper-responsive to setmelanotide, a melanocortin 4 receptor agonist. Br. J. Pharmacol. 173, 2614–2621 (2016).

41. Collet, T. H. et al. Evaluation of a melanocortin-4 receptor (MC4R) agonist (Setmelanotide) in MC4R deficiency. Mol. Metab. 6, 1321–1329 (2017).

42. Paxinos, G. & Franklin, K. B. J. The mouse brain in stereotaxic coordinates: hard cover edition. Acad. Press 2nd Editio, 360 (2001).

